# Ancient DNA reveals differences in behaviour and sociality between brown bears and extinct cave bears

**DOI:** 10.1101/056119

**Authors:** Gloria G. Fortes, Aurora Grandal-d'Anglade, Ben Kolbe, Daniel Fernandes, Ioanna N. Meleg, Ana García-Vázquez, Ana C. Pinto-Llona, Silviu Constantin, Trino J. de Torres, Jose E. Ortiz, Christine Frischauf, Gernot Rabeder, Michael Hofreiter, Axel Barlow

**Affiliations:** Institute for Biochemistry and Biology. University of Potsdam, 14476 Potsdam OT Golm Germany; Department of Biology and Evolution, University of Ferrara, I-44121 Ferrara Italy; Instituto Universitario de Xeoloxla, Universidade da Coruna,, 15081 A Coruna Spain; School of Archaeology and Earth Institute, University College Dublin, Dublin 4, Ireland; “Emil Racoviţă” Institute of Speleology, Department of Karst inventory and cave protection,, 050711 Bucharest, Romania; c/o J. Villarías Instituto de Historia, Consejo Superior de Investigaciones Científicas, 28037 Madrid, Spain; Depto. de Ingeniería Geológica y Minera, Universidad Politécnica de Madrid, 28003 Madrid, Spain; hInstitute of Palaeontology, University of Vienna, A-1090 Vienna, Austria; iDepartment of Biology, The University of York, YO10 5DD, UK

**Keywords:** *Ursus spelaeus*, *Ursus arctos*, ancient DNA, sociality, homing, extinction

## Abstract

Ancient DNA studies have revolutionised the study of extinct species and populations, providing insights on phylogeny, phylogeography, admixture and demographic history. However, inferences on behaviour and sociality have been far less frequent. Here, we investigate the complete mitochondrial genomes of extinct Late Pleistocene cave bears and middle Holocene brown bears that each inhabited multiple geographically proximate caves in northern Spain. In cave bears, we find that, although most caves were occupied simultaneously, each cave almost exclusively contains a unique lineage of closely related haplotypes. This remarkable pattern suggests extreme fidelity to their birth site in cave bears, best described as homing behaviour, and that cave bears formed stable maternal social groups at least for hibernation. In contrast, brown bears do not show any strong association of mitochondrial lineage and cave, suggesting that these two closely related species differed in aspects of their behaviour and sociality. This difference is likely to have contributed to cave bear extinction, which occurred at a time in which competition for caves between bears and humans was likely intense and the ability to rapidly colonise new hibernation sites would have been crucial for the survival of a species so dependent on caves for hibernation as cave bears. Our study demonstrates the potential of ancient DNA to uncover patterns of behaviour and sociality in ancient species and populations, even those that went extinct many tens of thousands of years ago.

## INTRODUCTION

Behaviour and sociality represent key mechanisms allowing populations to rapidly adapt to changing environments, to better exploit available resources, and also to resist pressures such as predation or climatic extremes that may negatively affect survival probability. Conversely, some behaviours could be maladaptive in certain contexts, particularly when populations are exposed to new and/or rapidly changing selective pressures, and may ultimately lead to population or even species extinction. Ancient animal remains can hold information on their behaviour and sociality. Spatial and temporal patterns of association among individuals can be investigated using standard paleontological and isotopic methods, and their relatedness can-at least in principle-be determined using ancient DNA approaches. The later, however, may represent a considerable technical challenge, as advanced DNA degradation will complicate recovery of suitable data that allows fine-scale resolution of genetic relationships among sufficient numbers of individuals to achieve statistical power.

Bears that lived in Eurasia during the Pleistocene represent a group that may be amenable to behavioural investigations using ancient DNA. Two major species (or species complexes) were widespread and sympatric in Pleistocene Eurasia: brown bears (*Ursus arctos*), that survived through the last glacial maximum (LGM) and are currently widespread across the entire Holarctic region; and the cave bear (*Ursus spelaeus* complex), an iconic representative of the Pleistocene megafauna, that went extinct prior to the LGM (Pacher & Stuart 2009; Stiller *et al.* 2010; 2014). For cave bears in particular, their habit to hibernate in caves has resulted in assemblages consisting of the bones of thousands of individuals at some sites, providing the opportunity to investigate uniquely well-defined fossil populations, deposited within an environment that enhances DNA preservation (Hofreiter *et al.* 2015). Although ancient brown bear remains typically occur at a much lower frequency in caves in comparison to cave bears, comprehensive palaeontological surveys of some caves have produced sufficient samples for population-level analysis (e.g. in Kurten 1968).

The factors that drove the cave bear to extinction have been subject to considerable study and discussion (Kurten 1968, Grayson & Delpech 2003; Pacher & Stuart 2009; Stiller *et al.* 2010). In agreement with palaeontological data, genetic studies of cave bears have found high genetic diversity and a large and constant population size until 50,000 yBP, followed by a decrease until its ultimate extinction around 24,000 yBP (Pacher & Stuart 2009; Stiller *et al.* 2010; 2014). Thus, the onset of decline of cave bear populations would have started around 25,000 years before the LGM, and is therefore not associated with any periods of substantial climatic change in Europe (Stiller *et al.* 2010; 2014). Brown bears, in contrast, show no evidence of population size changes coinciding with the cave bear population decline (Stiller *et al.* 2010). It has been argued that human activities played a major role in cave bear extinction (Grayson & Delpech 2003; Knapp *et al.* 2009; Munzel & Conard 2004; Bon *et al.* 2011; Stiller *et al.* 2014). However, explanations of why human activities could have so profoundly affected cave bear populations and not brown bear populations remain elusive. Differences in behaviour between the two species may have played a role, but identifying such differences is challenging because many aspects of cave bear behaviour remain uncertain. For example, paleontological studies of some cave bear caves have identified multiple depressions (hibernation beds or *bauges*, as described by Koby in 1953) in the cave floor that are thought to have been formed by hibernating bears. While this suggests communal hibernation, it is uncertain whether these were social or even family groups, or rather random assemblages of individuals forced together through competition for hibernation sites. Although genetic data could allow testing of such hypotheses, only a few studies have examined the population structure of cave bears at a local-i.e. individual cave-scale (Orlando *et al.* 2002; Richards *et al.* 2008; Hofreiter *et al.* 2004; Bon *et al.* 2011). Moreover, these studies were all based on short mtDNA fragments, which does not allow fine scale resolution of the genetic relationship between individuals.

In this study, we investigate complete mitochondrial genome sequences generated from the subfossil remains of multiple cave bears and brown bears from several caves in the North of Spain (Fig. 1). Four of the cave bear caves are located in close proximity (within a radius of 10km) within the Serra do Courel mountains (NW Spain), while the fifth one is located 450 km away in Navarra (NE Spain). The brown bear caves are also in close proximity (within a radius of 50km). In all cases, there are no apparent topographic barriers separating caves from one another. Thus, for such large bodied and presumably highly mobile mammals as cave bears and brown bears, movement between these caves would, in general, not have represented any significant challenge. In cave bears, we find that, even though caves were occupied simultaneously, each cave almost exclusively contained a unique clade of closely related haplotypes. This remarkable pattern suggests that cave bears returned to the cave where they were born and formed stable maternal social groups for hibernation. In brown bears, however, no such pattern is found suggesting greater flexibility with regard to hibernation site in this closely related species. We discuss the implications of these behavioural differences for the extinction of the cave bear, in addition to the wider potential of ancient DNA for the study of behavioural ecology, sociality, and extinction.

**Figure 1.**
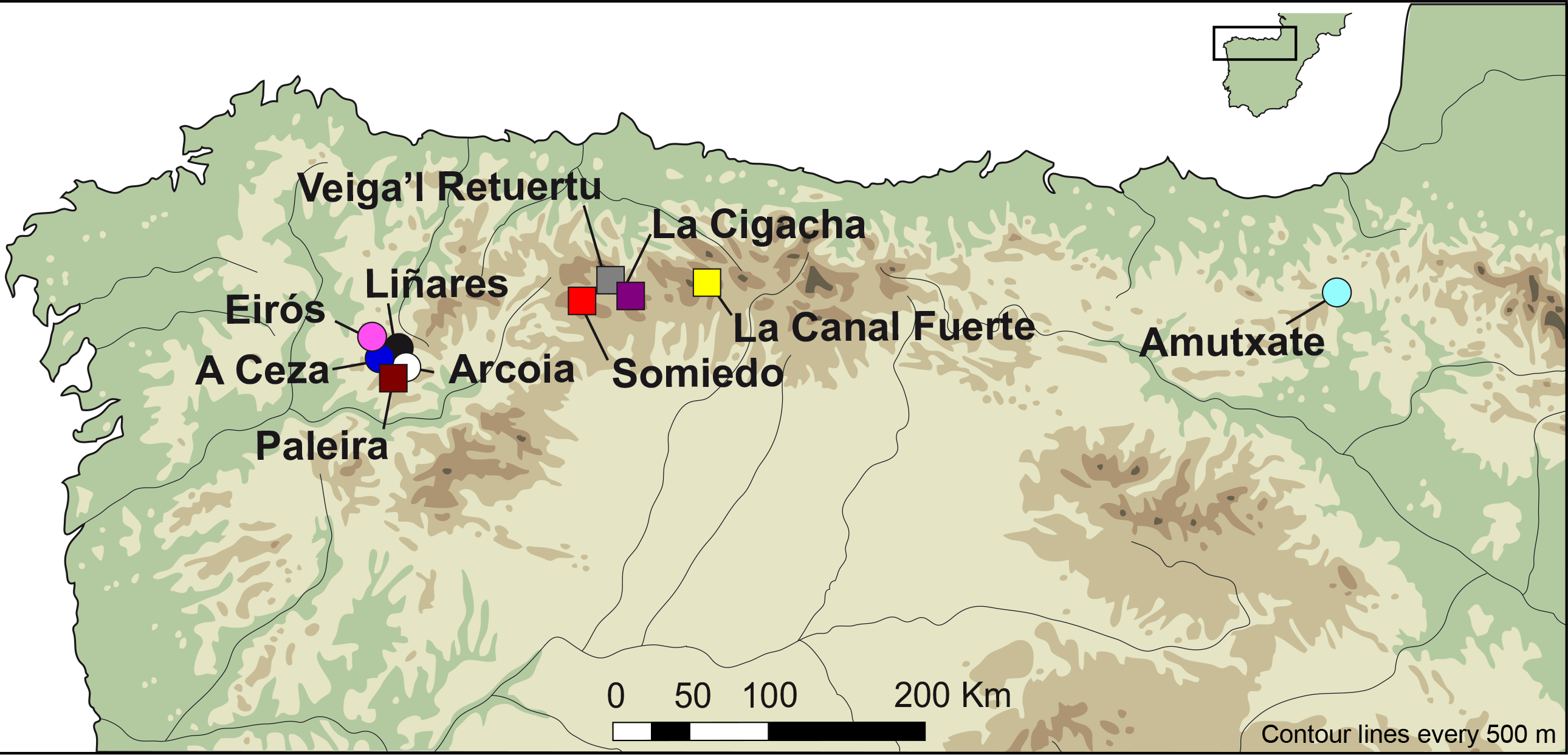
Map of Northern Spain showing locations of the caves investigated in this study. Circles represent sites with cave bears. Squares are sites with brown bears. Colours are consistent with Fig. 2.

**Figure 2.**
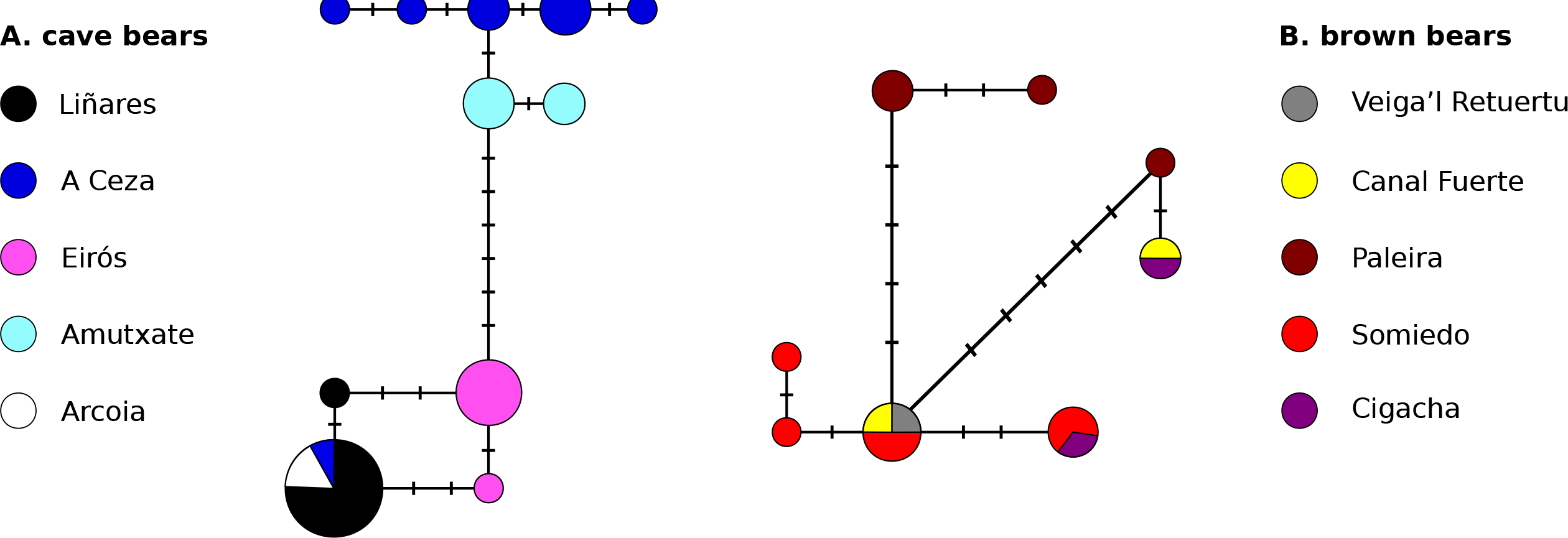
Haplotype networks of A. Iberian cave bears and B. Iberian brown bears, coloured according to the cave in which that haplotype was found (indicated next to each network). Circles are sized relative to haplotype frequency. Dashes along edges indicate single nucleotide mutations.

## MATERIALS AND METHODS

### Methods overview

We generated mitogenome sequences of cave bears and brown bears from their skeletal remains found in the caves shown in Figure 1. These sequences were used alongside published sequences obtained from GenBank to compare the maternal relatedness of individuals occurring within caves with that occurring among caves using haplotype network analysis, phylogenetic analysis and trait-phylogeny association tests. Finally, the ages of individuals were estimated using a combination of ^14^C and molecular dating. In particular, we investigated whether the occupation of caves was likely simultaneous, or instead temporally separated.

All but one of the novel Spanish bear mitogenome sequences reported here were obtained in a single experiment (we refer to as Experiment 1) that used hybridisation capture to enrich sequencing libraries for mtDNA prior to high-throughput sequencing. The details of Experiment 1 are reported below. A single Spanish cave bear sequence (sample E-VD-1838), in addition to sequences from seven bears from elsewhere in Europe, were obtained in separate experiments that are described in Section 1 of the Supporting Information.

### Sampling locations

The focal specimens used in this study were excavated in caves within karstic systems in the north-west of Spain, and were identified morphologically as either *U. spelaeus* or *U. arctos*. All of these sites represent natural accumulations and none of the remains are in archaeological context. Individual samples originated from different individual animals, identified based on age, sex or spatial distribution of the remains. Initially, specimens from 19 cave sites were investigated. These comprised 85 individuals from nine caves containing cave bear remains, and 24 individuals from ten caves containing brown bear remains. Many of these failed initial screening to identify samples that were likely permit recovery of the complete mitogenome sequence (see below), which limited sampling to five brown bear caves and five cave bear caves (shown in Fig. 1). Full details of the caves and samples investigated are provided in Section 2, Tables S1 & S2, and Fig. S1 of the Supporting Information.

### DNA extraction and sample screening

All pre-amplification aDNA analyses were performed in dedicated aDNA laboratories at the University of York (UK) or at the University of Potsdam (Germany). The compact part of bones, either femur, tibia, ribs, skull fragments or teeth, were utilised for DNA extraction. Prior to extraction, samples were UV irradiated for 10 minutes on each side and disposable cutting disks attached to a rotating electric drill were used to remove the outermost bone surface. For each sample, around 250 mg of cleaned bone was ground to powder using ceramic mortar and pestles. DNA extraction followed the protocol of Rohland *et al.* (2010).

DNA extracts were screened for likely presence and quality of endogenous DNA by attempting to PCR amplify 104bp and 126bp fragments of the mitochondrial control regions of cave bears and brown bears, respectively, using the primers described in Hofreiter *et al.* (2004) and a novel brown bear primer, UaF7 (5′-TCGTGCATTAATGGCGTG-3′). Amplification was assessed using agarose gel eletrophoresis and the authenticity of amplification products verified by Sanger sequencing, carried out in both directions using an ABI 3130XL at the Sequencing Service SAI (Servicios Centrais de Investigacion, University of A Coruña, Spain), followed by BLAST alignment of the consensus sequences.

### Sequencing library generation and hybridisation capture

We generated individually barcoded Illumina sequencing libraries using 20μl of those extracts for which short-amplicon PCR had previously been successful, following the protocol described in Meyer & Kircher (2010) with the following modifications. First, the filtration step between the blunt end repair and the adapter ligation was substituted by heat inactivation of the enzymes (Bollongino *et al.* 2013; Fortes and Paijmans 2015), in order to reduce the loss of short DNA fragments. Second, we used a double index barcoding system in which both the P5 and P7 adapters include a molecular barcode specific for each sample (Kircher *et al.* 2011; Fortes and Paijmans 2015). This facilitates the identification of chimeric molecules that could be formed during PCR amplification of the captured products. Library indexing and amplification involved 4 replicate parallel PCRs, each using 15 cycles, which were then pooled and purified using silica columns (Qiagen, France). The resulting cave bear and brown bear libraries were quantified using a Nanodrop Spectrophotometer (Thermo Scientific) and pooled, respectively, in equimolar quantities at a final concentration of 2 ng in 520 μl for hybridisation capture.

Hybridisation capture was carried out using 244k DNA SureSelect^TM^ microarrays (Agilent, Boblingen, Germany) with 2-fold tiling and 60bp probes. Separate arrays were used for the cave bear and brown bear library pools, with probes based on published mitogenome sequences of a Western European cave bear (EU327344, Bon *et al.* 2008) and brown bear (EU497665, Bon *et al.* 2008), respectively. Hybridisation capture followed the protocol of Hodges *et al.* (2009) with one modification. After the initial round of capture enrichment, library pools were amplified using primers IS5 and IS6 (Meyer & Kircher 2010) in 12 parallel PCRs and the resulting products were subjected to a second round of capture enrichment, as described in Fortes & Paijmans (2015).

### DNA sequencing and data processing

100bp single-end sequencing of mtDNA enriched library pools was carried out on a single lane of an Illumina HiSeq2000 instrument at the Danish National Sequencing Centre in the University of Copenhagen. The resulting BCL files were converted to fastq format using the Illumina base-calling pipeline (Illumina Pipeline v1.4). The program Cutadapt v1.3 (Martin, 2011) was then used to trim any P7 adapter sequences occurring at the 3' ends of reads, and a custom script used to identify and discard any reads that did not contain the appropriate P5 index, and then trim the index sequence from the remaining reads. Following this procedure, any reads < 25 bp were also discarded. The resulting cave bear and brown bear reads were then mapped to their respective reference mitogenome sequences used for capture probe design, using bwa-0.5.9 (Li & Durbin 2009) with seeding disabled, as suggested by Schubert *et al.* (2012). The alignment was sorted, filtered for minimum mapping quality (-q 30) and PCR duplicates removed using samtools (Li *et al.* 2009). The Mpileup tool in samtools 0.1.19-44428 was used to generate consensus sequences and to call polymorphic positions, using the-s option to specify a haploid genome. In order to prevent miscalling of polymorphic sites resulting from the presence of postmortem molecular damage to the ancient templates, the terminal five nucleotides at both 5' and 3' read ends were excluded from SNP calling, and for sites covered by less than 3 reads the bases were only called when all reads had the same nucleotide. All polymorphic sites identified in the vcf file were further checked by eye on Tablet version 1.13.05.02 (Milne *et al.* 2013). Read depth and coverage were determined using GATK (MacKenna *et al.* 2010). The presence of molecular damage characteristic of aDNA was confirmed using the software MapDamage (Ginolhac *et al.* 2011).

### Phylogenetic and network analysis

Only those novel sequences that provided > 70% total coverage of the mitogenome were used in subsequent analyses. Novel Spanish sequences were aligned along with seven novel sequences from ancient bears found elsewhere in Europe and 174 published mitogenome sequences from cave bears, brown bear and polar bears using the program MUSCLE (Edgar & Robert 2004) with default settings. A repetitive section of the d-loop was removed from the alignment as this was not recovered in many ancient samples and even when present could not be aligned unambiguously. All subsequent analyses used this alignment or subsamples of it.

To investigate the phylogenetic relation of Spanish cave bear and brown bear haplotypes to those occurring elsewhere in their respective distributions, we conducted phylogenetic analysis of the complete alignment under maximum likelihood (ML) using RAxML-HPC2 8.2.3 (Stamatakis, 2014) on the CIPRES Portal (Miller *et al.* 2010) using the American black bear (*U. americanus*) as outgroup. The ML tree was estimated under the GTR+G model and clade support assessed by 500 bootstrap replicates using the GTR+CAT model.

Networks of Spanish cave bear and brown bear haplotypes were then generated using the median-joining algorithm implemented in the program NETWORK (fluxus-engineering.com, Bandelt *et al.* 1999). To avoid any confounding effects of missing data on haplotype identification, all alignment columns containing missing data and/or alignment gaps were removed for network analysis.

We then investigated the strength of association of mitochondrial lineage and cave using trait-phylogeny association tests that account for phylogenetic uncertainty in the software BaTS (Parker *et al.* 2008). If mitochondrial phylogeny and cave are strongly associated, then the inferred number of changes in cave occupation across the phylogeny should be fewer than for a random prediction with no such association. We generated a Bayesian posterior sample of trees in BEAST v. 1.8.2 (Drummond *et al.* 2012), and then randomised the assignment of individuals to caves in order to generate a null distribution of the number of changes in cave occupancy when phylogeny and cave show no association. This strength of association was then tested by comparing this null distribution to the observed number of changes occurring across the posterior sample of trees using the parsimony score (PS) statistic (Slatkin & Maddison 1989). PS is a discrete metric and therefore models changes in cave occupation occurring across the phylogeny as discrete events.

To generate the posterior sample of trees used in trait-phylogeny association tests, the program PartitionFinder (Lanfear *et al.* 2012) was first used to select appropriate partitions and substitution models within each alignment (details in Section 2 of the Supporting Information, results in Tables S5 & S6, Supporting Information). BEAST analyses involved a coalescent Bayesian Skyline population model with unlinked substitution and strict clock models for each partition. Non-zero variation in substitution rates was rejected by preliminary runs using relaxed clock models. No clock calibrations were applied, and instead the substitution rate of the fastest-evolving partition was fixed to 1 and substitution rates for the remaining partitions estimated relative to the latter partition within open uniform priors between 0-2. MCMC chains ran for sufficient length to achieve convergence and sufficient sampling of all parameters (ESS > 200) after removal of burn-in, as verified in the program TRACER (Rambaut *et al.* 2014). LOGCOMBINER was used to remove pre-burn-in trees prior to trait-phylogeny association tests.

### Dating of cave lineages

Thirty-nine samples were directly ^14^C dated and 2-sigma calibrated using OxCal 4.2 online (accession date: 07/07/2015), based on the IntCal-13 curve (Reimer *et al.* 2013). For samples that lacked ^14^C dates, or were beyond the range of ^14^C dating, we estimated their ages using a Bayesian phylogenetic approach in BEAST (Shapiro *et al.* 2011). Phylogenetic age estimation was conducted individually for each undated cave bear and brown bear based on ^14^C dated representatives of their respective clades. We additionally tested the reliability of this procedure using a crossvalidation method, in which the age of each ^14^C dated sample was estimated and compared to its original ^14^C age. Due to the large number of individual analyses required, we a custom Perl script was used to automate the generation of BEAST input files. In each analysis, the posterior distribution of the tip date of the undated sample was sampled within an open uniform prior between 0 (present day) and one million years, both of which represent implausible extremes for the ages of these samples, while fixing the ages of ^14^C dated samples to the mean calibrated date. Substitution rates for all partitions were estimated within open uniform priors between 0-5x10^-7^ substitutions site^-1^ year^-1^. Other details of the BEAST analyses were as described above. Finally, we generated fully sampled calibrated phylogenies of the cave bear and brown bear clades by fixing tip dates to either mean calibrated ^14^C ages or median phylogenetic age estimates.

## RESULTS

### DNA sequences

PCR screening resulted in successful amplification of mitochondrial control region fragments in 57 out of 85 cave bear extracts and 23 out of 24 brown bear DNA extracts (details in Table S2, Supporting Information), which were then subjected to hybridisation capture enrichment and high-throughput sequencing. Mapping of sequence reads to their respective reference mitogenome sequences resulted in consensus sequences of 26 cave bears and 15 brown bears that were > 70% complete and used for further analysis (details in Table S4, Supporting Information). All datasets showed molecular damage patterns characteristic of ancient DNA (Figs. S2 & S3, Supporting Information). For cave bears, we added the sequence from an additional shotgun-sequenced individual (Section 1, Supporting Information) and previously published sequences from four other individuals from the focal caves, bringing the total number of Spanish cave bears analysed to 31.

Phylogenetic analysis supported the inclusion of these Spanish cave bear and brown bear sequences within the Western European *U. spelaeus* cave bear clade and the Western European brown bear clade 1 (Fig. S4, Supporting Information), identified by previous phylogeographic studies (Hirata *et al.* 2013; Stiller *et al.* 2014). Spanish cave bear and brown bear haplotypes were unique compared to all previously published haplotypes of conspecific bears occurring elsewhere in their respective distributions.

### Association of mitochondrial DNA and cave

Network analysis of Spanish cave bear haplotypes revealed close relationships between haplotypes found within the same cave (Fig. 2a). Most caves contain multiple unique haplotypes that are separated from each other by single nucleotide mutations. For example, Eirós and Amutxate caves each contain two unique haplotypes differing from one another by a single nucleotide mutation. Similarly, five unique and closely related haplotypes were found in A Ceza cave, but with the addition of a more divergent haplotype found in a single A Ceza individual (sample C7) that is shared with individuals from Arcoia and Liñares. An additional unique haplotype was found in Liñares cave that differs from this shared haplotype by a single nucleotide mutation. Even considering the occurrence of a single haplotype that is shared among three caves, an overall pattern of separation of haplotype clusters into caves is clear and obvious. Trait-phylogeny association tests further confirmed this pattern, showing fewer observed changes in cave occupation than expected by random (observed mean 5.9, null mean 18.0, p < 0.001), indicating a strong association of Spanish cave bear mitochondrial lineages with particular caves.

In contrast, an obvious segregation of mitochondrial haplotypes among different caves was not observed in middle Holocene Spanish brown bears (Fig. 2b). Haplotypes are widely shared among caves, with the exception of Pena Paleira, which contains three unique haplotypes, but these are not closely related. Trait-phylogeny association tests found the observed number of changes in cave occupation to not differ significantly from random (observed mean 6.5, null mean 8.2, p = 0.08), indicating a lack of statistically significant association between mitochondrial lineage and cave in these middle Holocene Spanish brown bears.

The association of mitochondrial haplotype lineage and cave revealed by network analysis for Iberian cave bears, but not for Iberian Holocene brown bears, is also evident from the time-calibrated phylogenies of their respective clades (Figs. 3 & 4). In addition, the broader geographic sampling of cave bear haplotypes in this analysis reveals that Spanish haplotypes as a whole are not monophyletic, with some cave linages sharing more recent common ancestry with haplotypes found in France and/or Germany.

**Figure 3.**
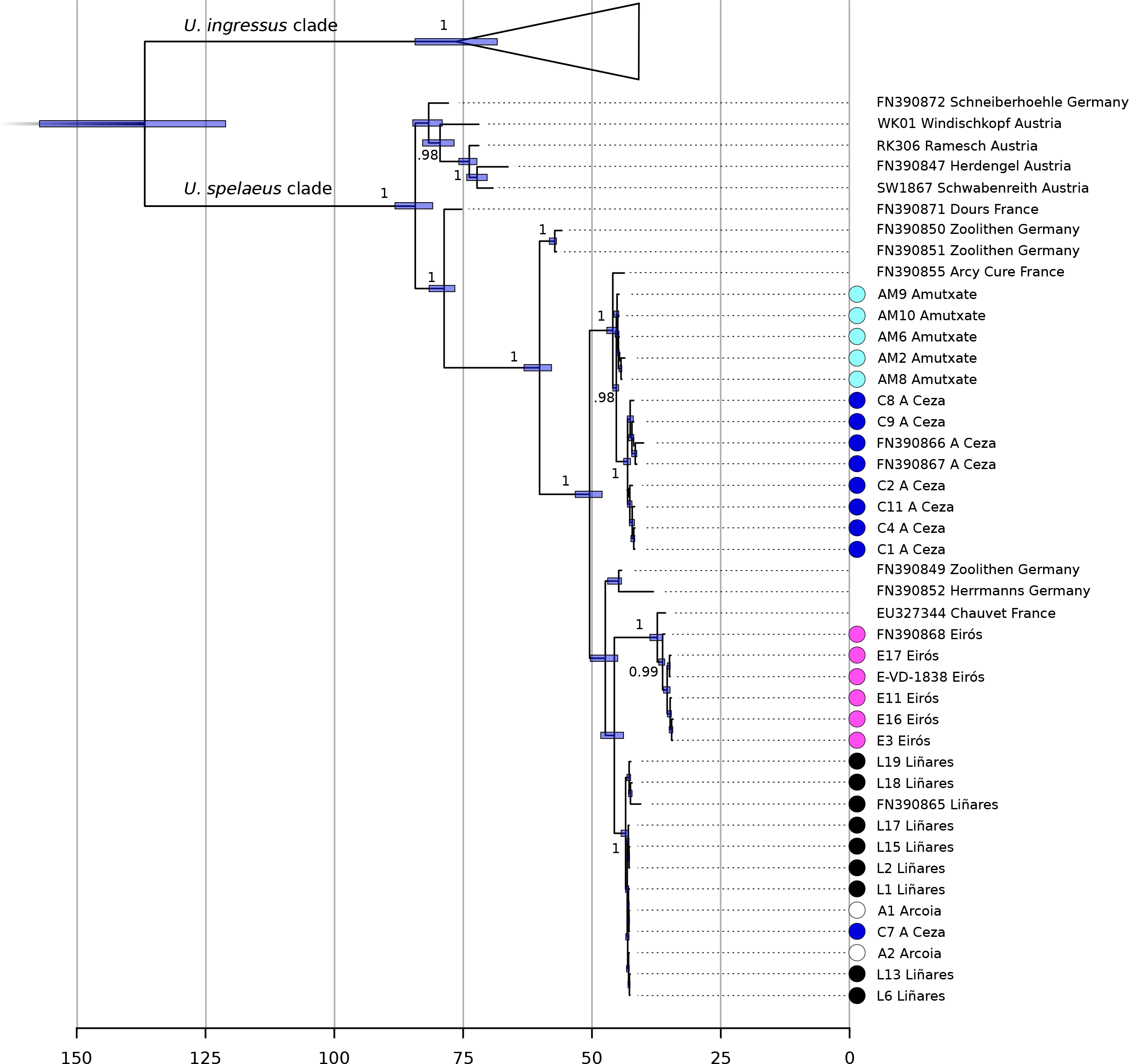
Time calibrated phylogeny of the Western European U. spelaeus cave bear clade. The lower scale shows kyBP. Branch labels indicate posterior clade probabilities ≥0.95, except for terminal tip clades where labels have been removed for simplicity. Nodes are centered on the median estimated divergence time and bars show the 95% HPD. Circles next to taxon names indicate Iberian cave bears and are coloured according to cave (consistent with Fig. 2). The *U. ingressus* clade that is sister to the *U. spelaeus* clade and was utilised for molecular dating is shown collapsed for simplicity.

**Figure 4.**
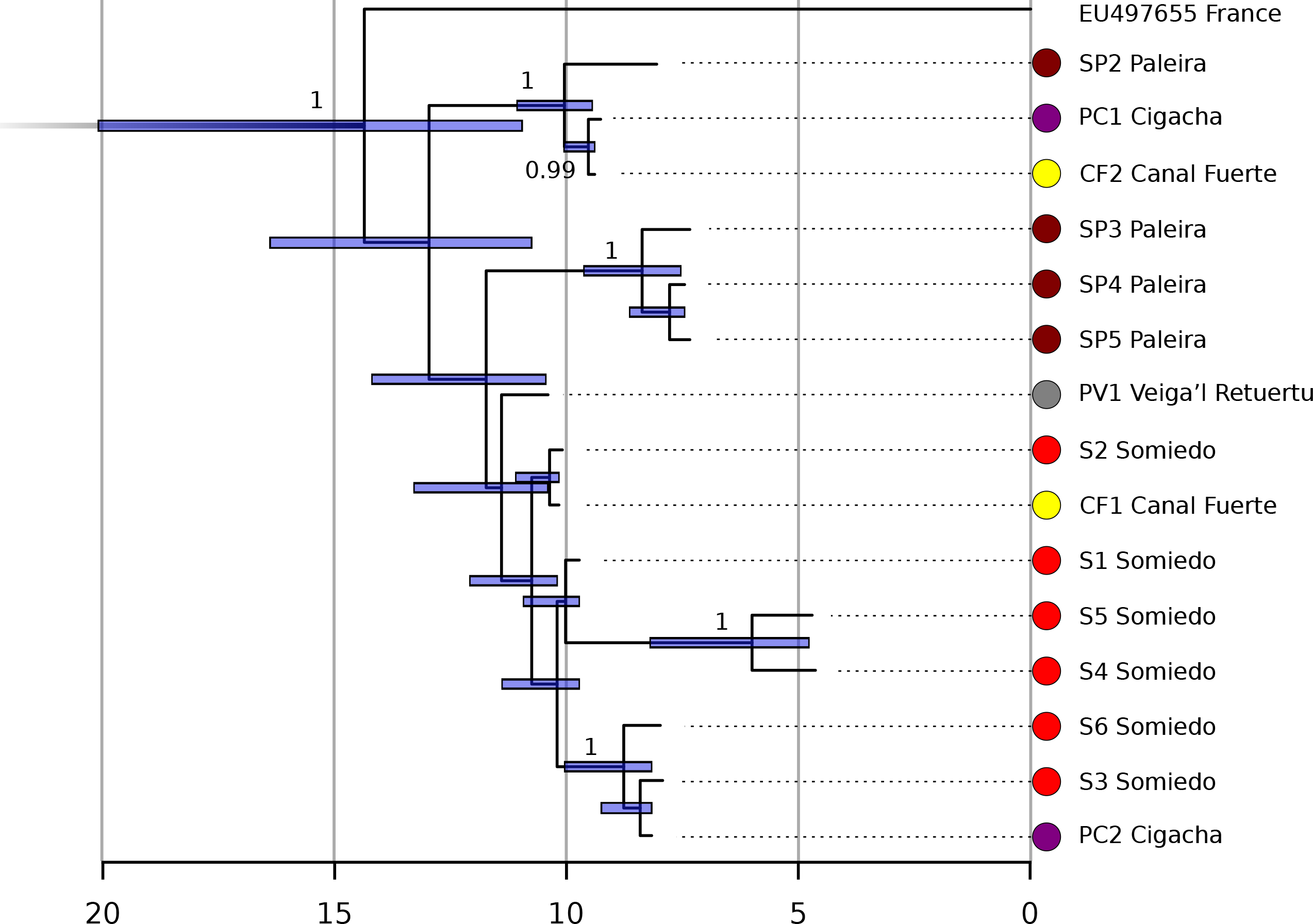
Time calibrated phylogeny of the Western European brown bear clade. The lower scale shows kyBP. Branch labels indicate posterior clade probabilities ≥0.95. Circles next to taxon names indicate Iberian brown bears and are coloured according to cave (consistent with Fig. 2). Two additional representatives of the West European brown bear clade, from Austria (sample Uap) and Bulgaria (GenBank Accession AP012591), were analysed and found to form a well supported sister lineage to the clade shown here that diverged an estimated 68,401 yBP ago (95% HPD 50,409.92,631 yBP). This lineage is not shown in order to better visualise divergence times
among Iberian brown bear haplotypes.

### Dating

^14^C ages spanned a range of > 40,000 to 28,251 yBP for cave bears and 41,201 to 2,520 yBP for brown bears (Table S3, Supporting Information).

Crossvalidation testing of the phylogenetic age estimation procedure resulted in 95% highest posterior densities (HPDs) that included the actual ^14^C age for all brown bears and all but one cave bear. Median estimated ages were also very close to the known age in most cases (Figs. S5 & S6, Supporting Information). These results support the reliability of this approach in estimating the ages of samples without ^14^C dates. Furthermore, age estimation for undated samples produced unimodal posterior estimates that are consistent with other sources of age information, where available, such as samples that were outside the range of ^14^C dating and those dated by amino acid racemisation (Table S7, Supporting Information).

Age estimates for cave bears (Fig. 5a) are compatible with the contemporaneous existence of the A Ceza, Amutxate, Arcoia and Liñares mitochondrial lineages. Although phylogenetic age estimates are associated with substantial uncertainty, the 95% HPDs of age estimates for these four caves show considerable overlap and median estimated ages are broadly comparable with each other, and with ^14^C dated samples. The simultaneous occupation of these caves is also supported by ^14^C dating of other specimens not included in this study (Pérez-Rama *et al.* 2011). In contrast to these caves, the Eiros mitochondrial lineage appears to have existed more recently and potentially without temporal overlap with those from other caves, although we do find slight overlap of Eiros ^14^C dates and HPDs from other caves in some cases (Fig. 5b). Generally younger ^14^C dates of Eirós in comparison to the other caves have also been reported previously, however, a single specimen was dated to more than 40,000 yBP (Pérez-Rama *et al.* 2011), and may therefore have existed contemporaneously with individuals from other caves. Unfortunately, this sample failed to yield any usable DNA and so its phylogenetic relation to more recent Eiros cave bears remains unknown. Caves containing brown bear remains were almost certainly inhabited simultaneously. ^14^C ages and a single phylogenetic estimate indicate temporal overlap in the habitation of these five caves between approximately 10,000 and 6,500 yBP (Fig. 5b).

**Figure 5.**
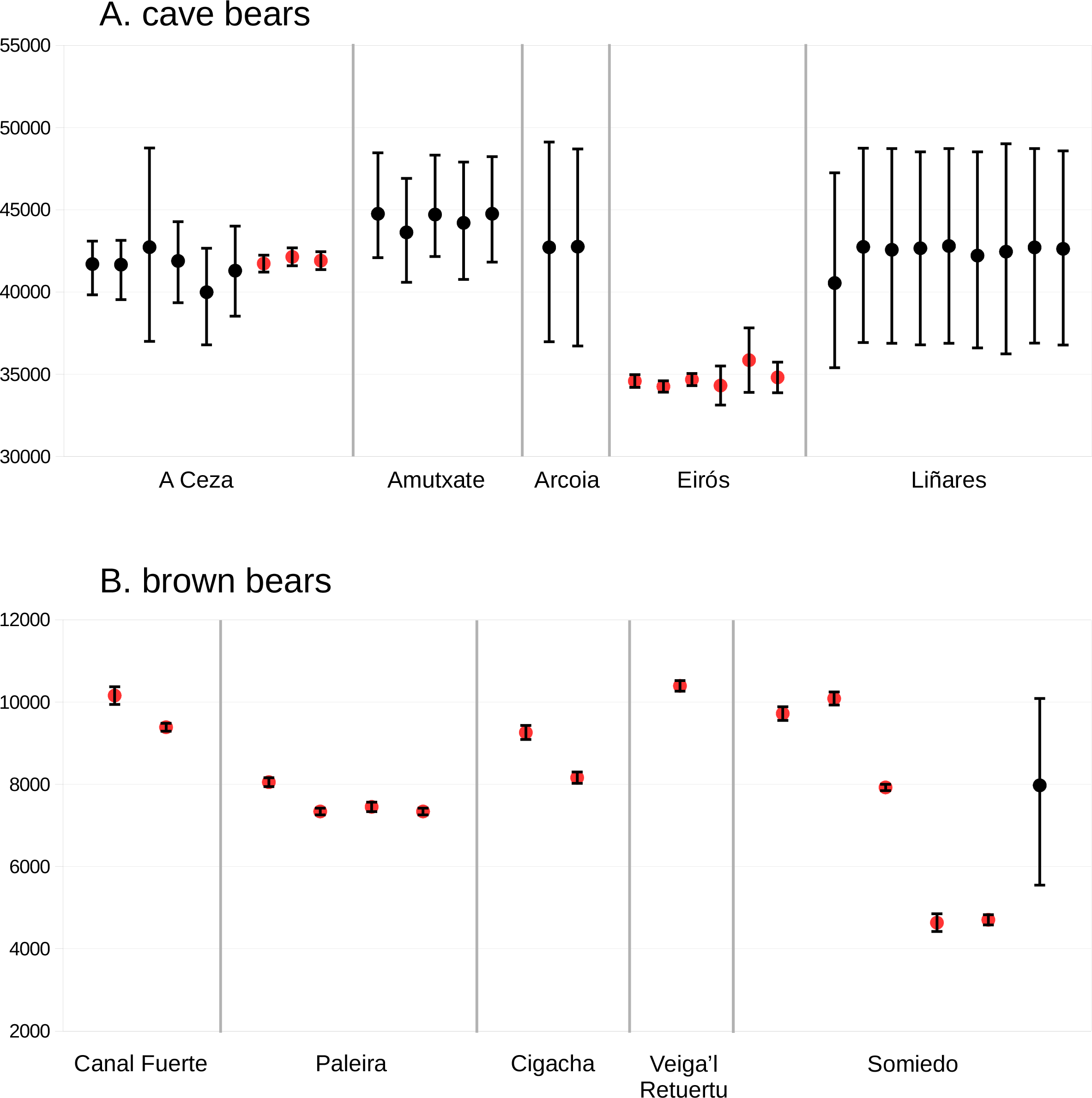
Time lines of A. Iberian cave bear and B. Iberian brown bear sample ages. Time in yBP is shown on the Y axes. Each point indicates the estimated age of an individual bear. Black points are median phylogenetic age estimates and red points are mean calibrated ^14^C ages. Error bars show 95% HPD and calibrated ^14^C uncertainty for phylogenetic age estimates and ^14^C ages, respectively.

## DISCUSSION

### Evidence for homing behaviour

Cave bears and brown bears that died in caves in the north of Spain show remarkably contrasting patterns of mitochondrial haplotype segregation. While no significant association of mitochondrial haplotypes and cave is found in middle Holocene brown bears, in the case of Late Pleistocene cave bears each cave contains, almost exclusively, a unique clade of closely related haplotypes. This structure exists despite caves being located in close geographic proximity and being inhabited simultaneously. We therefore interpret this as evidence of homing behaviour in cave bears. This scenario would involve a single intermixing cave bear population within which individuals-both males and females-returned to their native caves annually for hibernation, that is, the cave in which their mother hibernated and also gave birth, as demonstrated by the large amounts of perinatal individuals in the sites (Torres *et al.* 2002; Pérez-Rama *et al.* 2011).Such homing behaviour does not exclude mating between bears from different caves, but would have sorted the mitochondrial lineages by caves. In contrast, the lack of association between mitochondrial haplotype and cave in middle Holocene brown bears rejects this type of homing behaviour in this closely related species. This is further supported by studies of extant brown bear populations which show greater flexibility with regard to hibernation site than inferred here for cave bears (e.g. in Naves & Palomero 1993).

Evidence suggests that cave bears hibernated communally (e.g. Philippe & Fosse 2003). Homing behaviour would therefore result in non-random groups of close maternal relatives assembled at each cave. Thus, this behaviour can be further considered as a form of sociality. The temporal stability of these social groups is demonstrated by the observation of multiple unique haplotypes within caves that differ from their nearest relative by a single nucleotide substitution (Fig. 2). This suggests that within-cave haplotype variability is the result of nucleotide mutations that occurred during the period of cave occupation, most likely over thousands of years. A stepwise pattern of haplotype variability within caves has previously been reported for short cave bear control region sequences from the Ach valley, south-western Germany (Hofreiter *et al.* 2007), which in light of our finding suggests the potential for similar homing behaviour in that population. The temporal stability of cave occupation by cave bears is further demonstrated by two morphologically distinct cave bear forms that each occupied separate caves located only a few kilometers apart in Austria. These morphotypes sort into respective, genetically divergent mitochondrial clades. Despite their close proximity, a previous study found no evidence of haplotype exchange between caves even though simultaneous occupation over thousands of years, implying both site fidelity and reproductive isolation (Hofreiter *et al.* 2004). In the case of Spanish cave bears, however, we consider reproductive isolation unlikely due to a lack of any obvious morphological separation and relatively low levels of haplotype divergence between caves. Our preferred alternative, a single population with homing behaviour, makes specific predictions about patterns of nuclear DNA divergence among caves, and obtaining such data would be a valuable direction for future cave bear research.

Although we found a clear association of mitochondrial lineage and cave in Spanish cave bears, the association is not perfect. Specifically, we found a single haplotype that is shared among three caves: Liñares, A Ceza and Arcoia. This shared haplotype is common among Liñares individuals, and separated from a second Liñares haplotype by a single nucleotide mutation. In the second cave, A Ceza, the shared haplotype is considerably diverged from other haplotypes within that cave. In the third cave, Arcoia, both samples investigated have the shared haplotype. These later samples are the remains of juvenile individuals and no other cave bear remains have been found in this cave, raising the possibility that these juveniles (and potentially the A Ceza individual carrying the same haplotype) originate from Liñares. Regardless of the origin of this shared haplotype, while this pattern does imply some degree of movement between caves, the overall evidence for homing behaviour is clear and substantial. An ability to disperse and occupy other caves is further indicated by the sister group relationship found between Eirós cave haplotypes and a haplotype from Chauvet cave in France, two caves that were occupied simultaneously (see Table S3, Supporting Information; Bon *et al.* 2008; 2011). Thus, the Eirós haplotype lineage may be the result of long distance dispersal by female bears from distant caves, rather than movement among localised Spanish caves, which is also consistent with the apparent temporal separation of this lineage from the other Spanish caves.

### Wider implications

Homing behaviour has wider implications for species survival and conservation. For example, in extant black bears (*Ursus americanus*), it has been discussed as a potential problem for repopulation programs, as both females and males are able to track back to their home area after being captured by humans and released several kilometres away (Beeman & Pelton, 1976; Rogers & Lynn 1986; Clark *et al.* 2002). The same effect has been observed in Asian black bears (*Ursus thibetanus*), where genetic studies showed that 63% of the translocated bears migrate back to their original sites (Mukesh *et al.* 2015). Other well known examples include anadromous fishes, whose ability to return to breeding sites is affected by anthropogenic disruption of freshwater river systems (e.g. Pess *et al.* 2014), and similarly in marine turtles, where anthropogenic coastal development threatens habitats used for egg deposition (e.g. Wallace *et al.* 2011). Although ancient DNA provides the potential to investigate such behavioural patterns in species that have already gone extinct, behavioural inferences based on ancient DNA have been rare (notable examples are Huynen *et al.* 2010; and Allentoft *et al.* 2015). Our study clearly demonstrates the potential utility of ancient DNA in the study of behavioural ecology by revealing evidence of homing behaviour in extinct cave bears, and furthermore, through comparison with a closely related extant species, we have also uncovered clues on the potential causes of cave bear extinction.

The role of humans in the extinction of the cave bear has been debated (Grayson & Delpech 2003; Munzel & Conrad 2004; Knapp *et al.* 2009; Bon *et al.* 2011; Stiller *et al.* 2014), but explanations that also account for the survival of the sympatric brown bear have remained elusive. It is likely that the high dependence of cave bears on their native caves would have made them more sensitive to human competition for caves for several reasons. First, as noted previously (Grayson *et al.* 2003; Stiller *et al.* 2010), the generally high dependence of cave bears on caves for hibernation would have brought them into severe competition with humans (both Neanderthals and modern humans). Second, their tendency to come back to the same cave site would have made them comparatively predictable prey, which fits to the growing evidence of cave bear hunting, again by both Neanderthals and modern humans (Munzel & Conrad 2004; Wojtal *et al.* 2015). And third, this homing behaviour would have prevented a rapid recolonisation of empty caves from neighbouring populations. Overall, these factors could have contributed to the extinction of the cave bear as modern human populations expanded from Eastern to Western Europe, indeed, advancing in the same direction as the subsequent cave bear extinction. This is in agreement with recent studies that have questioned the relative contribution of Pleistocene climatic changes to cave bear extinction, and suggested instead a major impact of human activities (Knapp *et al.* 2009; Bon *et al.* 2011; Stiller *et al.* 2014). Finally, the lack of evidence of homing behaviour to their maternal caves in Spanish brown bears, a species that lived in widespread sympatry with cave bears but survived the human expansion into Western Europe, further implicates this behaviour as a factor in the extinction of the cave bear.

## ACKNOWLEDGMENTS

This work was supported by Xunta de Galicia, Conselleria de Economia e Industria, (Grant number 10 PXIB 162 125 PR to GGF); Ministerio de Economía y Competitividad (MINECO CGL2014-57209-P to AGD); European Science Foundation Research Networking Programmes (ConGenOmics, Ref. 5882 to GGF); ERC consolidator grant GeneFlow (310763 to MH); and KARSTHIVES Project funded by CNCS-UEFISCDI (PCCE_ID_31/2010 to SC). We also thank Andrea Manica for useful comments on the manuscript, Dr. Marius Robu, from the “Emil Racoviţă” Institute of Speleology for providing the sample PA1 from Romania, and Stefanie Hartmann for bioinformatic assistance.

## DATA ACCESSIBILITY

DNA sequences from the cave and brown bears obtained in this study are deposited in Genbank with accession numbers: XXXXX. DNA sequence alignment has been deposited in Dryad with accession number XXXXXX

## AUTHORS CONTRIBUTIONS

G.G.F, A.B, A.G and M.H designed and conceived of the study; G.G.F, A.B and I.N.M. performed molecular work; G.G.F, A.B, B.K and D.F, performed NGS data processing and statistical analysis; A.G, A.G.V, A.C.P, S.C, T.J.T, J.E.O, C.F and G.R collected and identified the ancient remains. G.G.F, A.B, and M.H drafted the manuscript with input from A.G and A.G.V. All authors gave final approval for publication.

